# Interactions between high-intensity light and unrestricted vision in the drive for hyperopia

**DOI:** 10.1101/2024.06.11.598474

**Authors:** Sayantan Biswas, Joanna Marie Fianza Busoy, Veluchamy A. Barathi, Arumugam R. Muralidharan, Leopold Schmetterer, Biten K. Kathrani, Noel A. Brennan, Raymond P. Najjar

## Abstract

**PURPOSE:** To evaluate the impact of optical versus illuminance factors and their duration-dependency on lens-induced hyperopia (LIH) in chick eyes.

**METHODS:** Hyperopia was induced in one eye in chicks (10 groups, n=126) from day 1 (D1) post-hatching until D8 using +10 D lenses with fellow eyes as controls. One group (LIH) served as the control without any interventions. Remaining groups were exposed to 2, 4 or 6 hours of unrestricted vision (UnV), high intensity light (HL), or both (HL +UnV). Ocular axial length (AL), refractive error, and choroidal thickness were measured on days 1, 4, and 8. Inter-ocular difference (IOD = experimental - contralateral eye) ± SEM was used to express outcome measures.

**RESULTS:** By D8, LIH decreased AL (-0.42±0.03 mm) and produced hyperopic refraction (+3.48±0.32 D) and choroidal thickening (+85.81±35.23 µm) in the LIH group (all, P<0.001). Exposure to UnV reduced LIH (i.e., hyperopic refraction, axial shortening, and choroidal thickening) in a duration-dependent manner, whereas HL potentiated the development of LIH in a duration-dependent manner. When combined, UnV overpowered HL, with resultant impact on refraction and AL being close to UnV alone, except after 6 hours, when HL+UnV induced shorter AL compared to UnV alone (P=0.03).

**CONCLUSION:** Daily exposure to HL, UnV, and HL+UnV altered LIH in a duration-dependent manner with UnV and LIH producing competing signals. The signal generated by UnV was generally stronger than HL in combined exposure, yet longer durations of HL affected the drive for emmetropization in eyes with UnV.

## Introduction

Emmetropization is a visually guided phenomenon, aiming to optimally focus the image on the retina throughout the development of the eye.^1^ Experimental myopic or hyperopic defocus using positive or negative lenses in front of the eye respectively, degrades the quality of the retinal image, disrupts normal emmetropization, leads to abnormal ocular axial growth,^3,4^ and the development of refractive error.^2, 3^

The most common refractive error is myopia or near-sightedness. Myopia is a global epidemic with an exponential growth in its prevalence among children, adolescents, and young adults, especially in South and East Asia.^4^ In 2020, myopia affected nearly 30% of the world’s population and this burden is expected to rise to 50% by 2050.^5^ Poor vision associated with myopia poses a global public health issue as it not only impacts the quality of early life but also imposes socio-economic consequences and increases the risk of sight threatening conditions if left uncontrolled.^5^

Hyperopia is another type of refractive error characterized by hyperopic refraction and shorter axial length (AL) of the eye.^6^ It often starts at an early age and remains relatively stable throughout visual maturation.^7^ Both myopia and hyperopia can be induced in experimental animal models using negative or positive defocusing lenses.^8, 9^ The lenses degrade the quality of the retinal image, and lead to aberrant ocular axial growth change,^3,4^ and the development of refractive error.^2, 3^ Myopic defocusing lenses (i.e., positive powered lens) fitted in front of the eye in animal models result in lens-induced hyperopia (LIH) associated with decreased ocular elongation, hyperopic refraction, and thicker choroid.^8^ Besides inducing hyperopia as a condition, positive lenses convey a “STOP” signal to the eye.^9^ Study of this phenomenon may thus be useful in understanding and developing methods for controlling ocular growth, which may have application in the maintenance of hyperopic reserve, myopia prevention or slowing of myopic progression.^10^

Compared to the extensive research focusing on myopia development and progression, only a few studies have examined the development of hyperopia. In children, a transient thickening of the choroid is observed following 2 hours of myopic defocus (+3 D),^11^ while transient reduction in AL and associated choroidal thickening were observed in young adults with +3 D defocus within 15–60 minutes.^12, 13^ Hence, incorporation of lens-induced myopic defocus as an optical correction can potentially control ocular growth and retard myopia progression in children. Results of long-term myopic defocus in the form of under-correction of myopia, bifocals and progressive addition spectacles are not clinically promising. Nonetheless, contact lenses with plus power in the lens periphery, orthokeratology—which induces peripheral plus corneal power—and spectacles with positively powered lenslets all have been shown to slow myopic progression.^14^

Besides the optical “STOP” signal, there is a growing body of evidence showing a protective effect of increased light intensity on the development of myopia, axial elongation and choroidal thinning in animal^15-19^ and clinical studies alike.^20, 21^ Ashby et al^16^ assessed the influence of high-intensity light (HL) on LIH on young chicks and found HL to accelerate positive lens (+7 D) compensation, but the end point was the same as in the control light group (500 lux). Using dual powered lens (+10 D/-10 D), Zheng et al^22^ showed myopic defocus and HL to be additive against the myopiagenic hyperopic defocus.

Recently we have investigated the interactions between optical re-focus and HL in a lens-induced myopia model. Our findings suggest that HL (15,000 lux) and unrestricted vision (UnV) have an additive, duration-dependent effect, particularly when administered for 6 hours, on reducing the development of lens-induced myopia (LIM) in chickens.^17^ UnV for 2–6 hours was reported to reduce 37%–96% of LIM caused by hyperopic defocus.^17, 23^ In contrast, myopic defocus is less sensitive to UnV with only 9% reduction after 3 hours of UnV in chickens.^23^ Equally, 9 hours of UnV following 3 hours of myopic defocus resulted in significant hyperopic refraction.^23^ Even wearing a positive lens for 12 minutes per day and UnV for the remainder of time developed hyperopia and reduced ocular elongation in chickens.^24^ In summary, although the temporal relationship of refractive change, i.e., lens compensation to positive lens, is considered to be duration-dependent, it is non-linear.^25^ Findings from clinical studies suggest myopic defocus to be more enduring than hyperopic defocus, producing stronger compensatory signal and greater persistence of the effects of myopic defocus even after its cessation.^26^

To date, the duration-dependent and synergetic effect of HL and UnV is yet to be studied in an LIH animal model. In this study we explore the duration-dependent effect of (1) myopic defocus, (2) HL, (3) UnV and (4) their combinations on hyperopia development (i.e., the STOP signal for ocular growth).

## Methods

### Animals and experimental setup

The animals used in this study were treated in accordance with the Association for Research in Vision and Ophthalmology (ARVO) statement for the Use of Animals in Ophthalmic and Vision Research. The study protocol (IACUC 2019/SHS/1479) was approved by the Association for Assessment and Accreditation of Laboratory Animal Care International accredited Singapore Experimental Medicine Centre (SEMC) Institutional Animal Care and Use Committee.

A total of 126, one-day-old chicks (mixed Golden Comet/White Leghorn strain) were obtained from the National Large Animal Research facility and were randomly divided into 10 groups, with each group consisting of 11 to 13 animals. The chicks were raised for 9 days in a custom-built enclosure of 75-cm (length) × 55-cm (width) × 43-cm (height) designed to hold two high-intensity light-emitting diode (LED) light fixtures. Light-dark cycle of 12/12-hour from 7 am to 7 pm and the temperature (maintained between 28°C to 32°C) within the enclosure with food and water ad libitum. A HOBO Pendant data logger (UA-022-64; ONSET, Bourne, MA, USA) was used to monitor the light and temperature patterns. Square wave gratings of a repeated sequence of light and dark bars were fitted on the enclosure wall as accommodative cues. Depending on the location of the animal within the enclosure, the spatial frequency of the gratings ranged between 0.01 to 0.42 cycles/degree. To ensure that emmetropization in chicks is not affected by variations in accommodative responses,^27^ all experimental groups were exposed to an identical visual environment. On the final day 9 of the experiment, the chicks were administered a sedative mixture of 0.2 mL/kg ketamine and 0.1 mL/kg xylazine. Subsequently, they were euthanized by administering an overdose of sodium pentobarbitone directly to the heart.

### Background and Experimental light setup

Throughout the 12/12-hour light-dark cycle, all chicks were raised under background lighting conditions of 150 lux. To achieve this, six strips of ultra-bright LEDs (4000K, 2NFLS-NW LED; Super Bright LED, Inc, St. Louis, MO, USA) were securely positioned above the enclosure. For the HL group, four LED panels, each consisting of 64 LEDs, were used, providing an average of 15,000 lux when measured at chicken eye level for various gaze angles (up, down, left, right, front, back) within the enclosure. The lighting system was controlled by a programmable Helvar DIGIDIM 910 router (Helvar, Dartford Kent, UK). To ensure accuracy, light levels and spectra were assessed using a calibrated radiometer and spectroradiometer (ILT5000 and ILT950; International Light Technologies, Peabody, MA, USA).

### Hyperopia induction

Hyperopia was induced monocularly in all chicks from day 1 (D1) post-hatching until day 8 (D8). This was achieved by utilizing a customized convex defocusing lenses (La SER Eye Jewelry, Port St. Lucie, FL, USA) with a power of +10 ± 0.5 diopters (D). The lenses had a total diameter of 12.5 mm and an optic zone diameter of 10 mm, with a base curve of 6.68 mm. A three-dimensional printed lens holder, custom-designed for this purpose, was used to randomly fit the lens to one eye of each chick. To secure the positioning of the lenses on the chick’s eyes and facilitate removal during cleaning and light exposure (in some groups), the lens holders were attached to a separate base piece that was glued to the down surrounding the eye. Taking into consideration the 10 mm diameter of the optic zone, an estimated vertex distance of 3 mm (from the defocusing lens to the corneal apex), and a calculated distance of 4.49 mm from the posterior nodal point to the defocusing lens on D1 in chicks, the approximate open viewing visual angle was estimated to be around 76.5 degrees. However, it should be noted that the open viewing visual angle might have been underestimated as these calculations did not account for changes in pupil size.^28^ The lenses were worn for a duration of 8 days and were thoroughly cleaned three times per day to maintain their optical clarity. The fellow eye remained uncovered and served as a control within each individual animal.

### Experimental Groups

Monocular LIH was applied to all the 10 groups of chicks. Out of these, nine groups underwent various interventions, such as HL (15,000 Lux), UnV, or a combination of HL and UnV, each lasting for different durations (0, 2, 4, or 6 hours) centered at 12:00 pm. Further information regarding the experimental interventions can be found below and in the accompanying table 1.

**Table 1:** Details on experimental groups and interventions.

| Experimental group | Duration of intervention (Hours) | N | Experimental eye | Control eye | Experimental interventions |  |
| --- | --- | --- | --- | --- | --- | --- |
|  |  |  |  |  | High intensity light status (15,000 lux) | Lens status |
| **LIH** | 0 | 13 | +10D | No lens | Off | Not removed |
| **HL** | 2 | 13 | +10D | No lens | On | Not removed |
|  | 4 | 13 | +10D | No lens | On | Not removed |
|  | 6 | 13 | +10D | No lens | On | Not removed |
| **UnV** | 2 | 13 | +10D | No lens | Off | Removed |
|  | 4 | 13 | +10D | No lens | Off | Removed |
|  | 6 | 12 | +10D | No lens | Off | Removed |
| **HL + UnV** | 2 | 11 | +10D | No lens | On | Removed |
|  | 4 | 12 | +10D | No lens | On | Removed |
|  | 6 | 13 | +10D | No lens | On | Removed |

Abbreviations: LIH = lens-induced hyperopia, HL = high-intensity light, UnV = unrestricted vision.

### LIH group

A total of 13 chicks in this group were raised in background laboratory light conditions (150 lux), and they were not exposed to HL or UnV.

### High-Intensity Light Groups (LIH + HL)

All the 3 groups had 13 chicks each and were exposed to 2, 4, or 6 hours of HL (15,000 lux) every day without removal of the defocusing lenses and background light for the remainder of the light cycle.

### Unrestricted Vision Groups (LIH + UnV)

Defocusing lenses were removed for 2, 4, or 6 hours/day for the 3 groups (n = 13, 13, and 12). Only background light was used to raise during the light cycle throughout the experiment.

### High-Intensity Light and Unrestricted Vision Groups (LIH + HL + UnV)

The 3 groups (n = 11, 12, and 13) were exposed to 2, 4, or 6 hours of HL (15,000 lux) every day along without removal of the defocusing lenses. The groups were exposed to background light for the remainder of the light cycle.

### Ocular Measurements In Vivo

All ocular measurements were carried out in a dimly lit room (<5 lux) between 12PM and 5PM and the animals were randomly evaluated to reduce the impact of circadian rhythm on the outcome measures. The body weight, ocular AL, refractive error, choroidal thickness (CT), central corneal thickness (CCT), and anterior chamber depth (ACD) were measured in all animals on D1, day 4 (D4) and D8 following the protocol described elsewhere.^17, 29^ A few chicks (2–3 animals on D1) who would not keep the eyelid open needed lid retractor. The examiner carefully inserted the lid retractor without touching the cornea or obstructing the examination procedure.

### Axial length

VuMAX HD (Sonomed Escalon, New Hyde Park, NY, USA) A-scan ultrasonography was used to measure the AL as described by Najjar et al.^29^ In summary, AL was defined as the distance between the echo spike originating from the anterior surface of the cornea and most anterior spike originating from the retina at a probe frequency of 10 MHz. A median of 7–10 scans were recorded as an individual reading.

### Refraction

A calibrated automated infrared photo-retinoscope was used as previously described,^30^ to measure ocular refraction. The chicks were gently held on an adjustable platform placed about one meter away from the infrared photo-refractor. The positioning of the chick’s head was done with great care to ensure optimal focus on its eye and to detect the first Purkinje image. Pupil size was adjusted for each eye and the median of the most hyperopic refraction readings (i.e., resting refraction) without any accommodative changes was calculated from the continuous refraction trace comprising at least 300 readings over time in each eye.^17, 29^

### Choroidal Thickness and Anterior Segment

Posterior segment spectral-domain optical coherence tomography (SD-OCT; Spectralis; Heidelberg Engineering, Inc., Heidelberg, Germany) was used to measure CT, whereas anterior segment OCT (RTVue; Optovue, Inc., Fremont, CA, USA) was used to image the anterior segment (ACD and CCT) as per the protocols described in Najjar et al.^29^ For both the procedures, the OCT operator gently held the alert chick’s head and positioned it in alignment with the OCT camera lens, allowing the infrared laser beam to enter the eye precisely through the center of the pupil. The centration of the pupil was further refined the alignment of the pupil, with multiple OCT scans obtained. The centration was within ±100 µm from the horizontal line for posterior segment OCT measurements. CT was defined as the distance between the inner border of the sclera and the outer border of the retinal pigment epithelium. The distance between the central most posterior layer of the cornea and the central most anterior layer of the lens was defined as the ACD, whereas CCT was defined as the average of three thickness measurements of the central cornea. The first author (SB), who was kept blind to the eye (LIH or control) and the study group conditions (HL, UnV, HL + UnV) throughout the measurement sessions, performed all the measurements manually.

### Analyses and Statistics

The data are presented as the mean ± SEM of the interocular difference (IOD) between the experimental (LIH) and the control eye (uncovered); calculated as the LIH eye – control eye. This approach accounts for the inter-animal variations in outcome measures due to the mixed breed and large number of animals (n = 126 chicks) included in this study. For comparing IODs in refraction, AL, CT, ACD, and CCT, a two-way repeated-measures ANOVA was employed. The factors considered were day, group, and the interaction between group and day. In case where the omnibus test indicated a significant interaction effect between group and day, pairwise multiple comparisons were conducted using the Holm-Sidak method. A two-way ANOVA was performed to assess the interaction between the type of intervention (HL, UnV, HL + UnV) and its duration (0, 2, 4, and 6 hours) on the refraction, AL, and CT. If the omnibus test yielded statistical significance, pairwise multiple comparisons were conducted using the Holm-Sidak method. For all statistical tests, the significance level was set at α = 0.05, and Sidak correction was applied for post hoc pairwise comparisons.

## Results

### Ocular Changes Associated with LIH

The LIH eyes developed hyperopic shift in refractive error (refraction: +5.12 ± 0.24 D and +7.39 ± 0.36 D by D4 and D8, respectively), primarily within the initial 4 days of +10 D lens wear (IOD: +1.31 ± 0.29 D and +3.48 ± 0.32 D by D4 and D8, respectively), in comparison to the uncovered contralateral control eyes (refraction: +3.81 ± 0.29 D and +3.91 ± 0.13 D by D4 and D8, respectively). Simultaneously, there was a reduced axial elongation in the LIH eyes (IOD: -0.28 ± 0.04 mm and -0.42 ± 0.03 mm by D4 and D8, respectively) and an increase in CT (IOD: 84.85 ± 19.05 µm and 85.81 ± 35.23 µm by D4 and D8, respectively) compared to the control eyes (all *P* <0.001) (Figure 1, 2 and 3, Supplementary Table S1). There was no difference in the CCT and ACD between LIH and control eyes (Supplementary Figures 1, 2).

**Figure 1.**
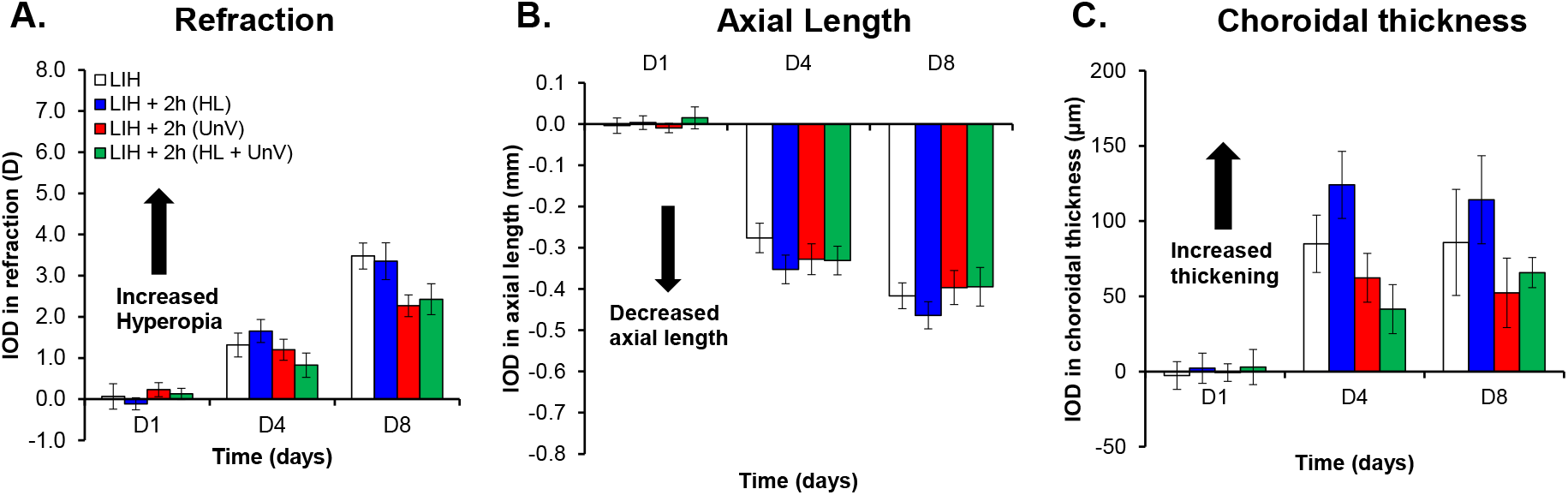
IOD in refraction, axial length, and choroidal thickness on days 1, 4, and 8 of the experimental protocol in the group not exposed to any intervention (LIH) and groups exposed to 2 hours of HL, UnV, or both (HL + UnV).

**Figure 2.**
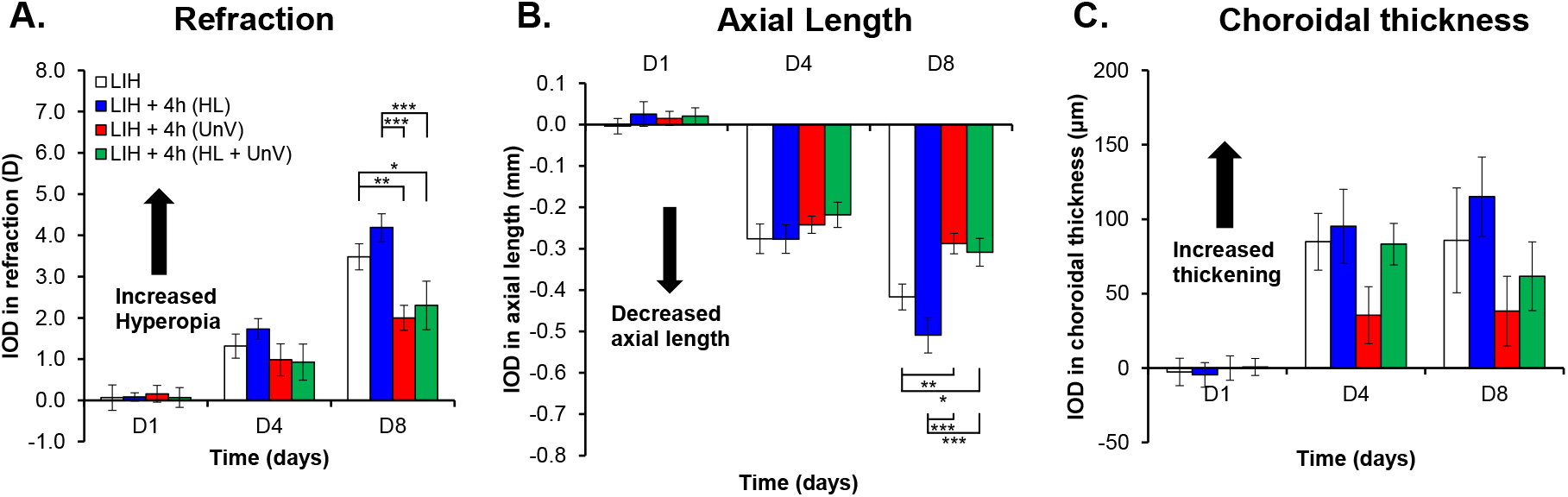
IOD in refraction, axial length, and choroidal thickness on days 1, 4, and 8 of the experimental protocol in the group not exposed to any intervention (LIH) and groups exposed to 4 hours of HL, UnV, or both (HL + UnV). For significant group effect: \**P <* 0.05, ^**^*P <* 0.01, ^***^*P <* 0.001 (two-way repeated-measures ANOVA).

**Figure 3.**
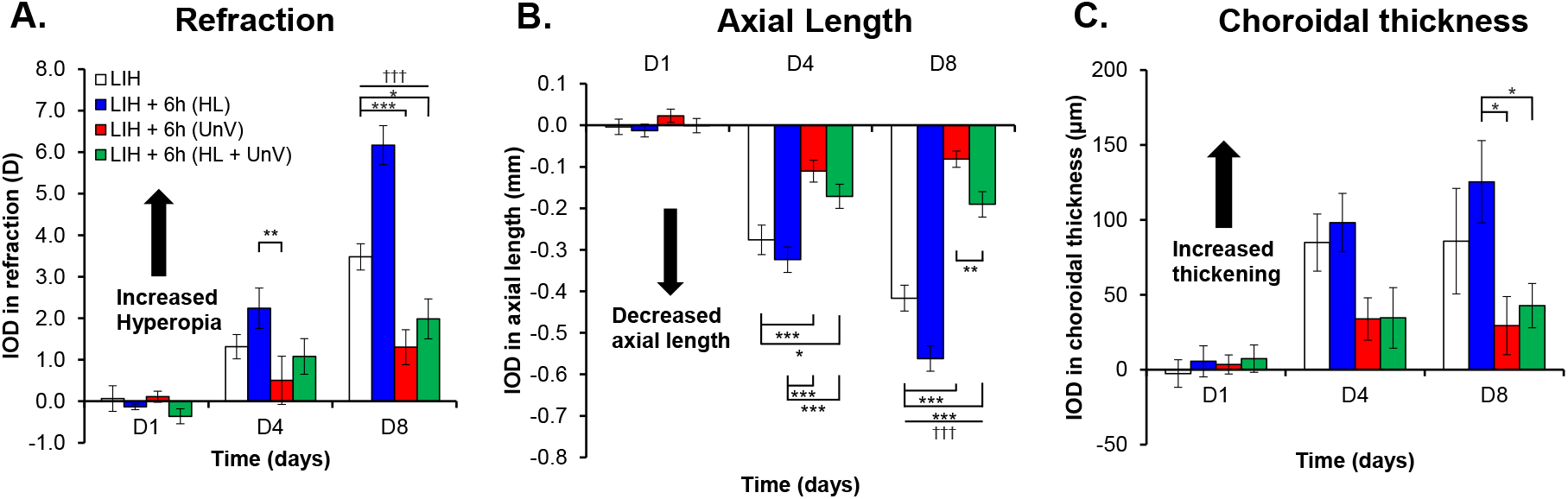
IOD in refraction, axial length, and choroidal thickness on days 1, 4, and 8 of the experimental protocol in the group not exposed to any intervention (LIH) and groups exposed to 6 hours of HL, UnV, or both (HL + UnV). All groups are significantly different from the LIH + HL group: ^†††^*P <* 0.001. For significant group effect: \**P <* 0.05, ^**^*P <* 0.01, ^***^*P <* 0.001 (two-way repeated-measures ANOVA).

### Impact of 2 hours of HL, UnV, and HL + UnV

For 2-hour interventions, IOD in refraction (F(2,46) = 82.53, *P* <0.001) (Figure 1A), AL (F(2,46) = 221.31, *P* <0.001) (Figure 1B) and CT (F(2,46) = 25.67, *P* <0.001) (Figure 1C) were only significantly different between the days of the intervention. Detailed results are available in Supplementary Table S1.

### Impact of 4 hours of HL, UnV, and HL + UnV

Four-hour interventions showed significant interactions between experimental group and day for IOD in refraction (F(6,92) = 2.38, *P* = 0.035). By D8, both 4 hours of UnV and HL+ UnV significantly reduced hyperopic refraction compared to the LIH group (both *P* < 0.05). UnV and HL+ UnV were equally effective (P >0.05) in reducing hyperopia. HL on the other hand significantly increased hyperopic refraction compared to both UnV and HL + UnV (both *P* <0.001) (Figure 2A). The group × day interaction was significant also for IOD in AL (F(6,94) = 4.59, *P* <0.001). Alike refraction, by D8, 4 hours of UnV and HL+ UnV significantly reduced axial elongation compared to the LIH group (both *P* <0.05). Equally HL was significantly effective in reducing axial elongation compared to both UnV and HL + UnV (both *P* <0.001) (Figure 2B). IOD in CT was only dependent on the day of the intervention (F(2,94) = 21.34, *P* <0.001, Figure 2C). Detailed results are available in Supplementary Table S1.

### Impact of 6 hours of HL, UnV, and HL + UnV

For 6-hour interventions, there was a significant group × day interaction for IOD in refraction (F(6,94) = 9.64, *P* <0.001). By D8, 6 hours of UnV (*P* <0.001) and HL + UnV (*P* = 0.011) significantly reduced hyperopic refraction, whereas HL alone increased hyperopic refraction compared to the LIH group (*P* <0.001). HL significantly increased hyperopic refraction compared to UnV on D4 and D8 (both *P* <0.01) and compared to HL + UnV on D8 (*P* <0.001) (Figure 3A). IOD in AL showed a significant group × day interaction (F(6,94) = 17.40, *P* <0.001), with UnV and HL + UnV showing increased axial elongation compared to the LIH eyes on D4 and D8 (LIH versus UnV: *P* <0.001 and LIH versus HL + UnV: *P* <0.05). On D8, HL produced significantly more reduction in AL compared to LIH (*P* <0.001). On both D4 and D8, HL exposed eyes had greater AL reduction than both UnV and HL + UnV (all *P* <0.001) (Figure 3B). IOD in CT was dependent on the group (F(3,94) = 4.04, *P* = 0.012) and day (F(2,94) = 17.61, *P* <0.001) individually, but their interactions did not reach statistical significance. CT in eyes exposed to HL were significantly higher than those with UnV (*P* = 0.024) or HL + UnV (*P* = 0.042) (Figure 3C). Detailed results are available in Supplementary Table S1.

### Impact of Experimental Interventions on ACD and CCT

IODs in ACD showed a significant effect of day for 2-hour (F(2,92) = 21.75, *P* <0.001), 4-hour (F(2,94) = 13.99, *P* <0.001), and 6-hour (F(2,94) = 20.34, *P* <0.001) interventions. IODs in CCT showed a significant effect of day only for 4-hour (F(2,94) = 4.12, *P* = 0.019), and 6-hour (F(2,94) = 7.39, *P* = 0.001) interventions (Supplementary Figures 1 and 2). For detailed results see supplementary table S1.

### Duration Response Curves on D4 and D8 of the Interventions

On D4, the impact of intervention on IODs in refraction (F(2,104) = 6.02, *P* = 0.003) was not duration dependent. For refraction, groups exposed to HL had significantly higher hyperopic refraction compared to those with UnV and HL + UnV (HL versus UnV: *P* = 0.008, HL versus HL + UnV: *P* = 0.009) (**Supplementary Figure 3A))**. The impact of the intervention on IODs of AL (F(2,104) = 14.15, *P* <0.001) was duration dependent. The interaction between the group and duration for IOD in AL was significant (F(4,104) = 2.98, *P* = 0.023), where 6 hours of HL was more effective in reducing ocular elongation than UnV (*P* <0.001) and HL + UnV (*P* = 0.001) (**Supplementary Figure 3B)**. IODs in CT were different between the intervention groups (F(2,104) = 9.36, *P* <0.001), with eyes exposed to HL having significantly thicker choroid than eyes exposed to UnV and HL + UnV (HL versus UnV: *P* <0.001, HL versus HL + UnV: *P* = 0.002) (**Supplementary Figure 3C)**.

On D8 of the protocol, there was a significant interaction between the duration and type of intervention on IODs of refraction (F(4,104) = 7.07, *P* <0.001). Both 4-hour and 6-hours of HL, but not 2-hours of HL, significantly increased hyperopic refraction induced by LIH compared to UnV (both 4 and 6-hour: *P* <0.001) and HL + UnV (4-hour: *P* = 0.003 and 6-hour: *P* <0.001) which decreased hyperopic refraction compared to LIH (4-hour: both UnV and HL + UnV: *P* <0.05; 6-hour: UnV: *P* <0.001 and HL + UnV: *P* = 0.011) (Figure 4A). Likewise, the interaction between the duration and type of intervention was significant for AL (F(4,104) = 9.87, *P* <0.001) where both 4-hour and 6-hours of HL, but not 2-hours of HL, further reduced AL compared to UnV (both 4 and 6-hour: *P* <0.001) and HL + UnV (both 4 and 6-hour: *P* <0.001) which increased AL compared to the LIH group (prevented AL shortening) (4-hour: both UnV and HL + UnV: *P* <0.05; 6-hour: both UnV and HL + UnV: *P* <0.001). For the 6-hour group, experimental eyes exposed to HL +UnV had shorter AL compared to eyes exposed to UnV (*P* = 0.028) (Figure 4B). IODs in CT (F(2,104) = 9.75, *P* <0.001) was different between groups across the different durations of the interventions, with HL inducing further choroidal thickening compared to LIH and compared to UnV (*P* <0.001) and HL + UnV (*P* = 0.003) (Figure 4C).

**Figure 4.**
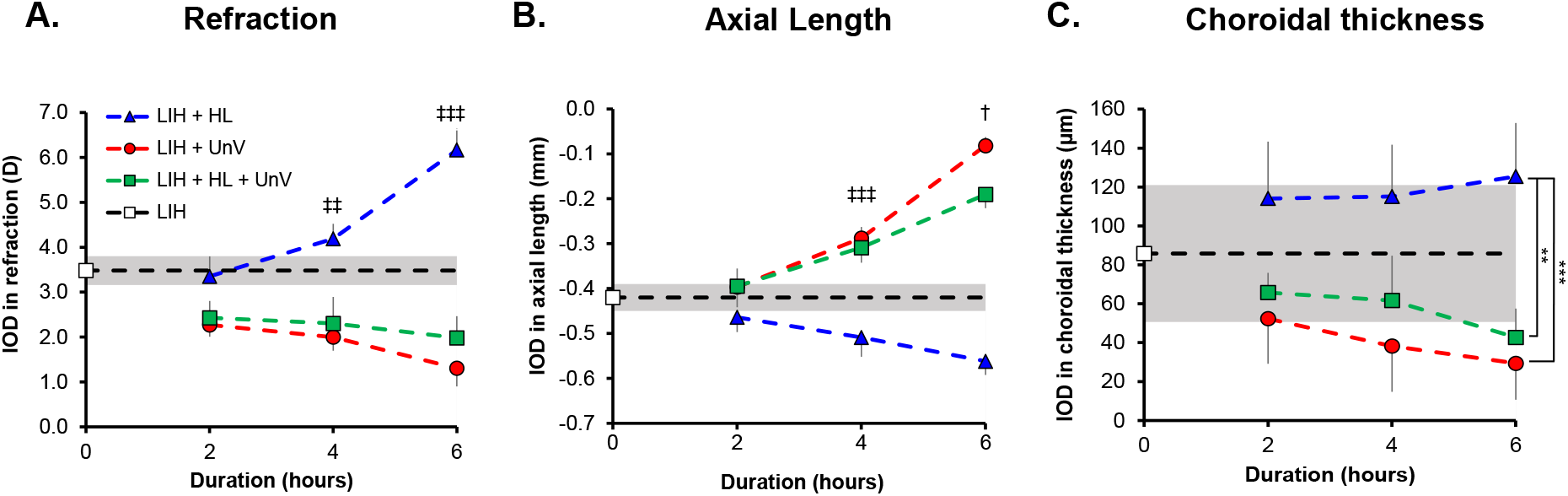
Duration-response curve for the IOD in refraction (**A**), axial length (**B**), and choroidal thickness (**C**) in the groups exposed to 2, 4, and 6 hours of HL, UnV, or both (HL + UnV) on day 8 of the experimental protocol. The LIH group that was not exposed to any intervention is represented by a *white square* and a *shaded area* for mean ± 95% confidence interval. HL is significantly different from the other two groups: ^‡‡^ *P <* 0.01, ^‡‡‡^*P* <0.001. All the groups are different from each other: ^†^*P <* 0.05 (at least). HL group is significantly different from both UnV and HL + UnV groups: ^*^P < 0.05, ^**^P < 0.01, ^***^P < 0.001.

## Discussion

In this study, we investigated the duration-dependent, differential, and combined effects of HL and UnV on the ocular growth STOP signal induced by LIH in a chicken model. The effect of HL, UnV, and HL + UnV in altering hyperopic refraction, AL elongation and CT were duration dependent by D8 of the intervention. Unlike in LIM, HL and UnV did not yield a similar effect in an LIH model. As previously reported,^17^ HL exacerbated the effects of LIH (i.e., increased hyperopic refraction, axial shortening and choroidal thickening) in a dose dependent matter, with the highest impact observed after 6 hours of exposure, followed by 4 and 2 hours. Conversely, UnV countered the effects of LIH (i.e., reduced hyperopic refraction, axial shortening and choroidal thickening) in a dose dependent manner with the highest being after 6 hours of exposure, followed by 4 and 2 hours. Interestingly, the impact of UnV overpowered HL with the combined effects of HL + UnV showing close similarity to UnV, except for AL after 6 hours of HL + UnV, where eyes exposed to LIH + HL + UnV had shorter ALs compared to eyes exposed to LIH + UnV alone. Consistent with previous findings, there was no significant change in ACD or CCT among the groups.^16^

The effect of UnV in reducing LIH in a duration-dependent manner has previously been reported by Schmid et al,^23^ where hyperopic refraction decreased by 8.4%, 27.7% and 42.2% on exposure to 3, 6 and 9 hours of UnV by D5, respectively. Correspondingly, exposure to UnV for 3, 6 and 9 hours per day increased AL elongation by 11.1%, 22.2% and 44.4%, respectively.^23^ In comparison, by D8 we observed 34.8%, 42.5%, 62.6% decrease in hyperopic refraction and 4.8%, 31%, 81% increase in AL elongation on exposure to 2, 4, and 6 hours of UnV, respectively. The increased impact of UnV observed in our study could potentially be attributed to disparities in the experimental protocol such as the age (visual maturation), strain of chickens, duration of the experimental protocol, as well as background lighting, visuo-spatial surroundings during UnV and the timing of UnV (centered around noon for this study and spilling into the afternoon). In fact, 2 hours of myopic defocus (+10 D) during noon or evening reduces ocular growth effectively as opposed to wearing +10 D lens continuously, whereas morning defocus induces less LIH. Similarly, 2 hours of positive lens removal in noon and evening caused increase in ocular growth more than morning removal.^31^ When it comes to the temporal dynamics of hyperopia induction, it has been proposed that temporal changes induced by compensation to positive lenses, although duration-dependent, is non-linear, as the rise and fall of the internal emmetropization signal is not directly proportional to the duration of lens wear, rather on the frequency of wear with short durations.^25^ In addition, earlier studies investigating the impact of UnV on LIH reported that interrupted hyperopia (UnV = 2 hours of relief from +4 D) resulted in a myopic shift in refractive state compared to the constant hyperopic group in tree shrews.^32^ These findings, along with ours, suggest that UnV pushes towards emmetropization based on the updated (i.e., the temporary hyperopic defocus created during UnV) state of image defocus. Conversely, using +5 D lens wear, Zhu and colleagues showed that even 30 minutes of UnV twice a day can result in a 43% increase in hyperopia in marmosets.^33^ These findings, although contradictory to ours, suggest that the inherent emmetropization signal to low myopic defocus (+5 D) does not decay when the treatment period is long (4 weeks) accompanied by multiple visual stimulation (UnV/ LIH × twice a day).

Exposing LIH eyes (+7 D) to HL (15,000 lux) for 5 hours per day, Ashby et al^16^ showed no change in refraction by D5 but a 46.2% reduction in axial elongation compared to LIH eyes without HL. In contrast, we recorded -3.7%, 20.4%, 77.3% increase in hyperopic refraction and 9.5%, 21.4%, 33.3% reduction in AL elongation relative to the contralateral control eye by D8 on exposure to 2, 4, and 6 hours of HL, respectively. In addition to the difference in experimental protocol, the experimental lights used by Ashby et al^16^ mimicked daylight (range 300-1000 nm, peak 700 nm), while our experimental lights had typical LED spectrum with two peaks around 449 nm and 583 nm. Recently a study on form deprivation myopia has shown the fullness of light spectrum may affect the refractive development in chicks.^34^

Nevertheless, our study agrees with Ashby et al’s^16^ findings at D4 on the concept that HL potentiates LIH and axial shortening, while UnV promotes emmetropization based on the updated ocular defocus status (i.e., the hyperopic eye without the positive lens), thus slowing LIH. Whether HL would still promote AL shortening had emmetropization been achieved (+10 D) is unclear. Yet, 6 hours of HL when combined with UnV triggered AL shortening compared to UnV alone (Figure 3B) thus suggesting that HL always promotes AL shortening rather than ocular compensation to defocus. These findings may explain a role of HL outdoors in protecting against myopia, through a potential build-up and maintenance of “hyperopic reserve” in growing eyes.

The choroid plays a role in the regulation of ocular growth and emmetropization. Choroidal thickening occurs in response to myopic defocus (positive lens).^35, 36^ Although studies on the effect of HL on CT under LIH are lacking, HL without LIH is expected to induce an increase in CT.^17, 34, 37^ Yet, we did not observe any increase in the CT of control eyes exposed to HL (i.e., HL, HL+UnV) compared to control eyes not exposure to HL (i.e., UnV). Conversely, HL in addition to positive lens, led to significantly thicker choroid compared to HL + UnV and UnV. This change in CT, is thought to be largely due to change in choroidal blood flow, permeability and vasodilation of choroidal vessels associated with the rise in intraocular temperature and neurotransmitter release.^38, 39^ By D8, LIH eyes exposed to 2, 4 and 6 hours of HL had choroidal thickening by 33%, 34.2% and 46.2% respectively, while eyes exposed to 2, 4 and 6 hours of HL + UnV and UnV had choroidal thinning by 23.4%, 28.1%, 50.3% and 39%, 55.5%, 65.8% respectively. Even though both HL + UnV and UnV resulted in decreased CT, HL + UnV, had slightly thicker choroid than UnV alone (P >0.05) (Figures 1-3C). Contrary to our finding, choroidal thickening by 16% was observed on removal of the myopic defocus (+5 D) for 30 minutes twice a day in marmoset eyes.^33^

HL and UnV probably trigger different mechanisms of action. UnV, being a visual/optical feedback guided phenomenon,^8, 40^ stops emmetropizing the eye at null IODs. In contrast, HL appears to work via a different pathway involving photoreceptor stimulation and releasing of retinal neurotransmitters.^15, 16, 41^ HL induced increase in retinal dopamine (DA) level is associated with lower LIM.^42^ However, the role of DA in positive lens compensation is unclear with mixed reports of both enhancement^43^ and no effect^44^ on LIH with injection of DA agonist such as apomorphine and 6-hydroxy DA, respectively. Studying the possible dopaminergic and cholinergic mechanisms of LIH development resulted in contradictory findings of increase,^45^ decrease or no change ^46, 47^ in retinal DA levels in eyes with LIH. Gamma-Aminobutyric acid (GABA) is another neurotransmitter related to the light exposure, is co-released alongside DA from the dopaminergic amacrine cells.^48^ Baclofen, a GABAB receptor agonist administration reduces LIH and CT, which further inhibits DA release and DOPAC content compared to LIH eyes without baclofen.^47^

Our study has a few limitations. First, it’s difficult to generalize our findings in chicks to humans given the differences between chicken and humans in their ocular anatomy and optics.^49^ The chicks were housed in a visual environment devoid of fine spatial details, color, and other regular features which promotes emmetropization.^50^ While the findings are in harmony with the literature suggesting that removing myopic defocus reduces hyperopia development, the finding is limited to animal models as humans are not subjected to myopic defocus in daily life. The other finding is that exposure to HL can potentiate hyperopia development in a duration-dependent manner regardless of the optical status of the eye. However, exposure to such high intensity (15,000 lux) of light for 16%, 33% or 50% (2, 4 or 6 hours) of the daytime is often not implementable in real life.

## Conclusion

In conclusion, our study showed that daily exposure to 2, 4, or 6 hours of UnV slows LIH by promoting emmetropization in a duration-dependent manner. The combination of UnV and HL of 2-4 hours does not potentiate the impact of UnV. Conversely, our findings suggest that HL potentiates the drive for hyperopia (slowing ocular growth) independent of the optical status of the eye. From a translational perspective, our findings also indirectly highlight the capability of long periods of exposure to HL to secure a hyperopic reserve in developing eyes, which may explain the protective effect of time outdoors against myopia onset.

## Acknowledgements

The authors thank Mr. Noel Sng for helping with the design of the chicken enclosure.

Supported by research grant from Singapore Government (IAF), Industry Collaboration Project Grant (I1901E0038), and Johnson & Johnson to R.P. Najjar. The funding organization has no role in the design, conduct, and interpretation of the research.

Disclosure: **S. Biswas**, None; **J.M.F. Busoy**, None; **V.A. Barathi**, None; **A.R. Muralidharan**, None; **L. Schmetterer**, None; **B.K. Kathrani**, Johnson and Johnson (E); and **N.A. Brennan**, Johnson and Johnson (E); **R.P. Najjar**, None.

