## Supplementary material for "Interactions between high-intensity light and unrestricted vision in the drive for hyperopia"

**Supplementary table 1:** Changes in ocular measurements in groups exposed to 0h, 2h, 4h and 6h of high intensity light, unrestricted vision, or both. Data represented as mean ± SEM of the interocular difference between experimental and control eyes.

| Ocular parameter | Duration of intervention (hours) | Protocol | Days | | | *P-values 2W RM ANOVA | | |
|---|---|---|---|---|---|---|---|---|
| | | | D1 | D4 | D8 | Group | Day | Group x Day |
| Refraction (D) | 0 | LIH | 0.07 ± 0.31 | 1.31 ± 0.29 | 3.48 ± 0.32 | - | | |
| Axial length (mm) | | | 0.00 ± 0.02 | -0.28 ± 0.04 | -0.42 ± 0.03 | | | |
| Choroidal thickness (µm) | | | -2.62 ± 9.22 | 84.85 ± 19.05 | 85.81 ± 35.23 | | | |
| ACD (µm) | | | -0.01 ± 0.01 | -0.07 ± 0.01 | -0.02 ± 0.03 | | | |
| CCT (µm) | | | 2.08 ± 1.05 | 3.42 ± 1.42 | 2.56 ± 1.63 | | | |
| Refraction (D) | 2 | LIH + HL | -0.11 ± 0.15 | 1.66 ± 0.28 | 3.35 ± 0.45 | 0.054 | <0.001 | 0.066 |
| | | LIH + UnV | 0.23 ± 0.17 | 1.20 ± 0.26 | 2.27 ± 0.26 | | | |
| | | LIH + HL + UnV | 0.13 ± 0.14 | 0.83 ± 0.30 | 2.43 ± 0.37 | | | |
| Axial length (mm) | | LIH + HL | 0.00 ± 0.02 | -0.35 ± 0.03 | -0.46 ± 0.03 | 0.557 | <0.001 | 0.584 |
| | | LIH + UnV | -0.01 ± 0.01 | -0.33 ± 0.04 | -0.40 ± 0.04 | | | |
| | | LIH + HL + UnV | 0.02 ± 0.03 | -0.33 ± 0.03 | -0.39 ± 0.05 | | | |
| Choroidal thickness (µm) | | LIH + HL | 2.19 ± 10.04 | 124.08 ± 22.24 | 114.12 ± 29.26 | 0.089 | <0.001 | 0.34 |
| | | LIH + UnV | -0.69 ± 5.77 | 62.31 ± 16.23 | 52.31 ± 23.14 | | | |
| | | LIH + HL + UnV | 2.95 ± 11.64 | 41.45 ± 16.32 | 65.77 ± 10.06 | | | |
| ACD (µm) | | LIH + HL | -0.01 ± 0.02 | -0.09 ± 0.02 | -0.09 ± 0.02 | 0.08 | <0.001 | 0.413 |
| | | LIH + UnV | -0.01 ± 0.01 | -0.10 ± 0.01 | -0.11 ± 0.03 | | | |
| | | LIH + HL + UnV | 0.00 ± 0.01 | -0.09 ± 0.02 | -0.08 ± 0.01 | | | |
| CCT (µm) | | LIH + HL | 2.31 ± 1.50 | 2.31 ± 0.87 | 3.46 ± 2.01 | 0.676 | 0.054 | 0.497 |
| | | LIH + UnV | -1.00 ± 0.78 | 4.06 ± 0.70 | 1.73 ± 1.39 | | | |
| | | LIH + HL + UnV | 0.86 ± 1.30 | 3.25 ± 0.82 | 1.11 ± 1.55 | | | |
| Refraction (D) | 4 | LIH + HL | 0.08 ± 0.10 | 1.73 ± 0.26 | 4.19 ± 0.34 | <0.001 | <0.001 | 0.035 |
| | | LIH + UnV | 0.16 ± 0.20 | 0.98 ± 0.39 | 2.00 ± 0.30 | | | |
| | | LIH + HL + UnV | 0.07 ± 0.24 | 0.93 ± 0.44 | 2.30 ± 0.59 | | | |
| Axial length (mm) | | LIH + HL | 0.03 ± 0.03 | -0.28 ± 0.03 | -0.51 ± 0.04 | 0.008 | <0.001 | <0.001 |
| | | LIH + UnV | 0.02 ± 0.02 | -0.24 ± 0.02 | -0.29 ± 0.02 | | | |
| | | LIH + HL + UnV | 0.02 ± 0.02 | -0.22 ± 0.03 | -0.31 ± 0.03 | | | |
| Choroidal thickness (µm) | | LIH + HL | -4.62 ± 8.23 | 95.19 ± 24.92 | 115.12 ± 26.59 | 0.109 | <0.001 | 0.387 |
| | | LIH + UnV | -0.04 ± 8.13 | 35.42 ± 19.15 | 38.23 ± 23.45 | | | |
| | | LIH + HL + UnV | 0.71 ± 5.67 | 83.25 ± 13.98 | 61.67 ± 23.04 | | | |
| ACD (µm) | | LIH + HL | -0.01 ± 0.01 | -0.11 ± 0.02 | -0.12 ± 0.06 | 0.115 | <0.001 | 0.507 |
| | | LIH + UnV | -0.01 ± 0.01 | -0.08 ± 0.01 | -0.05 ± 0.02 | | | |
| | | LIH + HL + UnV | -0.01 ± 0.01 | -0.10 ± 0.02 | -0.11 ± 0.03 | | | |
| CCT (µm) | | LIH + HL | -0.46 ± 0.84 | 4.23 ± 0.84 | 1.88 ± 1.63 | 0.773 | 0.019 | 0.695 |

| | | | | | | | | |
|---|---|---|---|---|---|---|---|---|
| | | LIH + UnV | 2.19 ± 0.82 | 2.87 ± 0.79 | 3.01 ± 1.04 | | | |
| | | LIH + HL + UnV | 0.58 ± 0.88 | 3.17 ± 0.71 | 2.65 ± 1.50 | | | |
| Refraction (D) | 6 | LIH + HL | -0.13 ± 0.07 | 2.24 ± 0.49 | 6.17 ± 0.47 | <0.001 | <0.001 | <0.001 |
| | | LIH + UnV | 0.11 ± 0.13 | 0.50 ± 0.58 | 1.30 ± 0.42 | | | |
| | | LIH + HL + UnV | -0.36 ± 0.18 | 1.08 ± 0.43 | 1.98 ± 0.48 | | | |
| Axial length (mm) | | LIH + HL | -0.01 ± 0.02 | -0.32 ± 0.03 | -0.56 ± 0.03 | <0.001 | <0.001 | <0.001 |
| | | LIH + UnV | 0.02 ± 0.02 | -0.11 ± 0.03 | -0.08 ± 0.02 | | | |
| | | LIH + HL + UnV | 0.00 ± 0.02 | -0.17 ± 0.03 | -0.19 ± 0.03 | | | |
| Choroidal thickness (µm) | | LIH + HL | 5.58 ± 10.38 | 98.12 ± 19.50 | 125.42 ± 27.49 | 0.012 | <0.001 | 0.096 |
| | | LIH + UnV | 3.50 ± 6.38 | 33.83 ± 14.16 | 29.38 ± 19.45 | | | |
| | | LIH + HL + UnV | 7.35 ± 9.08 | 34.54 ± 20.21 | 42.69 ± 14.79 | | | |
| ACD (µm) | | LIH + HL | -0.01 ± 0.01 | -0.10 ± 0.02 | -0.10 ± 0.03 | 0.14 | <0.001 | 0.48 |
| | | LIH + UnV | -0.01 ± 0.01 | -0.08 ± 0.01 | -0.06 ± 0.02 | | | |
| | | LIH + HL + UnV | -0.01 ± 0.01 | -0.09 ± 0.01 | -0.07 ± 0.03 | | | |
| CCT (µm) | | LIH + HL | -1.54 ± 1.25 | 3.88 ± 1.38 | 1.58 ± 1.82 | 0.654 | 0.001 | 0.762 |
| | | LIH + UnV | 0.25 ± 0.57 | 3.31 ± 1.22 | 2.58 ± 1.10 | | | |
| | | LIH + HL + UnV | 0.81 ± 0.69 | 3.83 ± 0.75 | 2.22 ± 1.30 | | | |

All values expressed as the mean inter-ocular difference between experimental and control eyes ± SEM. 2W RM ANOVA: Two-way repeated measures analysis of variance, LIH: Lens induced hyperopia, HL: High-intensity light, UnV: unrestricted vision, ACD: Anterior chamber depth, CCT: Central corneal thickness

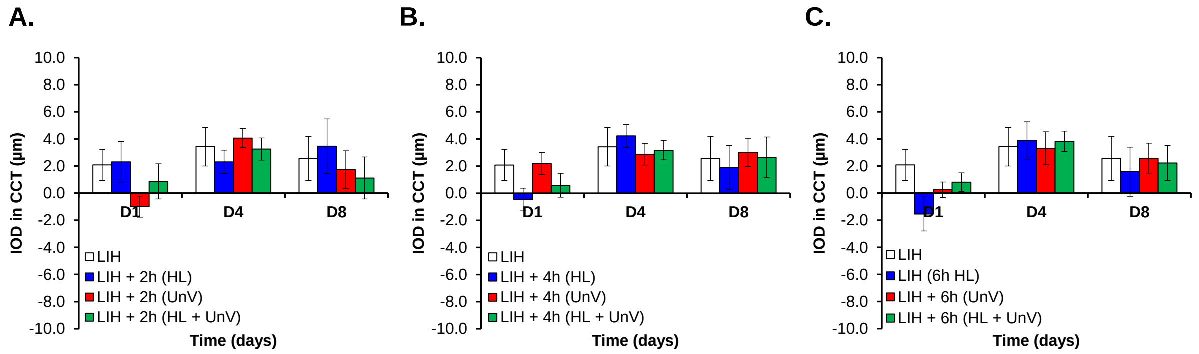

**Supplementary Figure 1.** IOD in CCT on days 1, 4, and 8 of the experimental protocol in the group not exposed to any intervention (LIH) and groups exposed to 2 hours (**A**), 4 hours (**B**), and 6 hours (**C**) of HL, UnV, or both (HL + UnV).

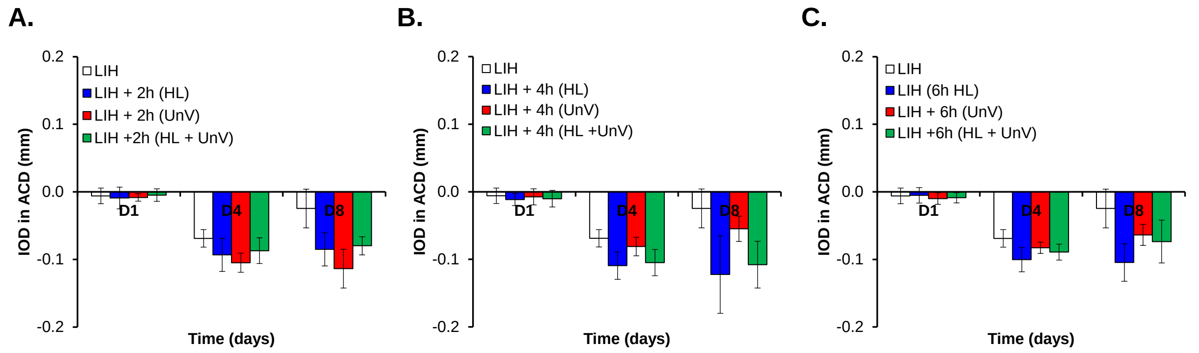

**Supplementary Figure 2.** IOD in ACD on days 1, 4, and 8 of the experimental protocol in the group not exposed to any intervention (LIH) and groups exposed to 2 hours (**A**), 4 hours (**B**), and 6 hours (**C**) of HL, UnV, or both (HL + UnV).

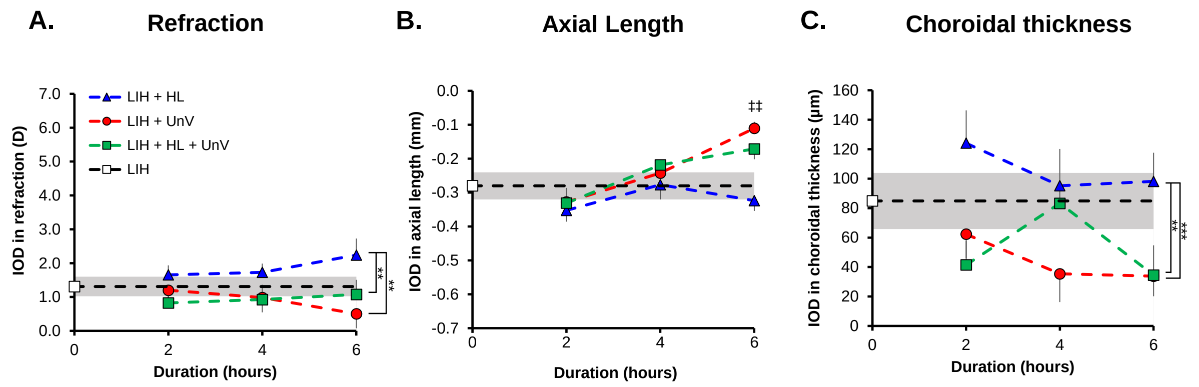

**Supplementary Figure 3.** Duration-response curve for the IOD in refraction (**A**), axial length (**B**), and choroidal thickness (**C**) in the groups exposed to 2, 4, and 6 hours of HL, UnV, or both (HL + UnV) on day 4 of the experimental protocol. The LIH group that was not exposed to any intervention is represented by a *white square* and a *shaded area* for mean ± 95% confidence interval. HL group is different from both UnV and HL + UnV groups at 6 hours: ‡‡P < 0.01. HL group is significantly different from both UnV and HL + UnV groups: **P* < 0.05, ***P* < 0.01, ***P < 0.001.
